## Supplementary information (figures, results, tables and methods) for "Intracortical recordings reveal Vision-to-Action cortical gradients driving human exogenous attention"

### Supplementary Results

*Clusters’ hemispheric lateralization*

To test if the clusters’ spatial distribution differs between right and left hemispheres, we performed a contingency table analysis only in symmetrically covered regions (309 contacts, 148 in the left hemisphere and 161 in the right hemisphere; see methods), that revealed a significant lateralization (*χ²*(5)=29.09, *p*<0.001). Post hoc comparisons showed that this effect resulted from a significant right lateralization of Cluster 2 and a significant left lateralization of Cluster 3 (post hoc binomial tests, *p*=0.01 and *p*=0.003).

### Supplementary Figures

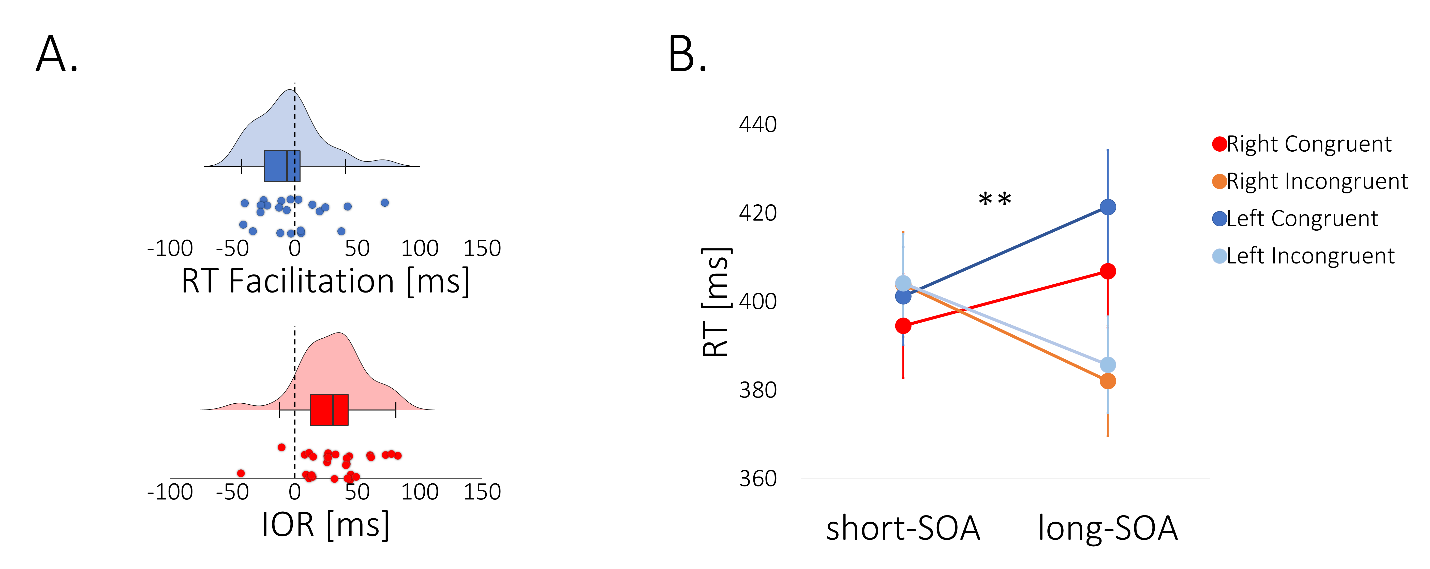

**Figure S1 -** Behavioral effects. (A) Individual RT effects. Raincloud plots of patient RT difference between Congruent and Incongruent trials, in the short-SOA condition (RT Facilitation effect; top; blue dots) and in the long-SOA condition (IOR effect; bottom; red dots). Shaded areas represent RT distributions for long-SOA (shaded red) and short-SOA (shaded blue) conditions; n=28 independent participants. (B) RT effects for right- & left-sided targets. Left target Congruent RTs were slower than Right target Congruent RTs, across both SOAs (repeated-measures 3-way ANOVA: Target-side X Congruence interaction - *F*_(1,27)_=8.28, *p*=0.008, *η^2^*=0.007, n=28 independent participants), reflecting the Poffenberger effect, i.e. faster RTs for right cue & target than for left cue & target, when responding with the right hand. In Incongruent trials in which cue & target appear at opposite sides of the screen, this effect might have averaged out. No other Target-side effects reached significance, and IOR and RT-facilitation effects did not significantly differ between left-sided and right-sided targets. ** *p*=0.008.

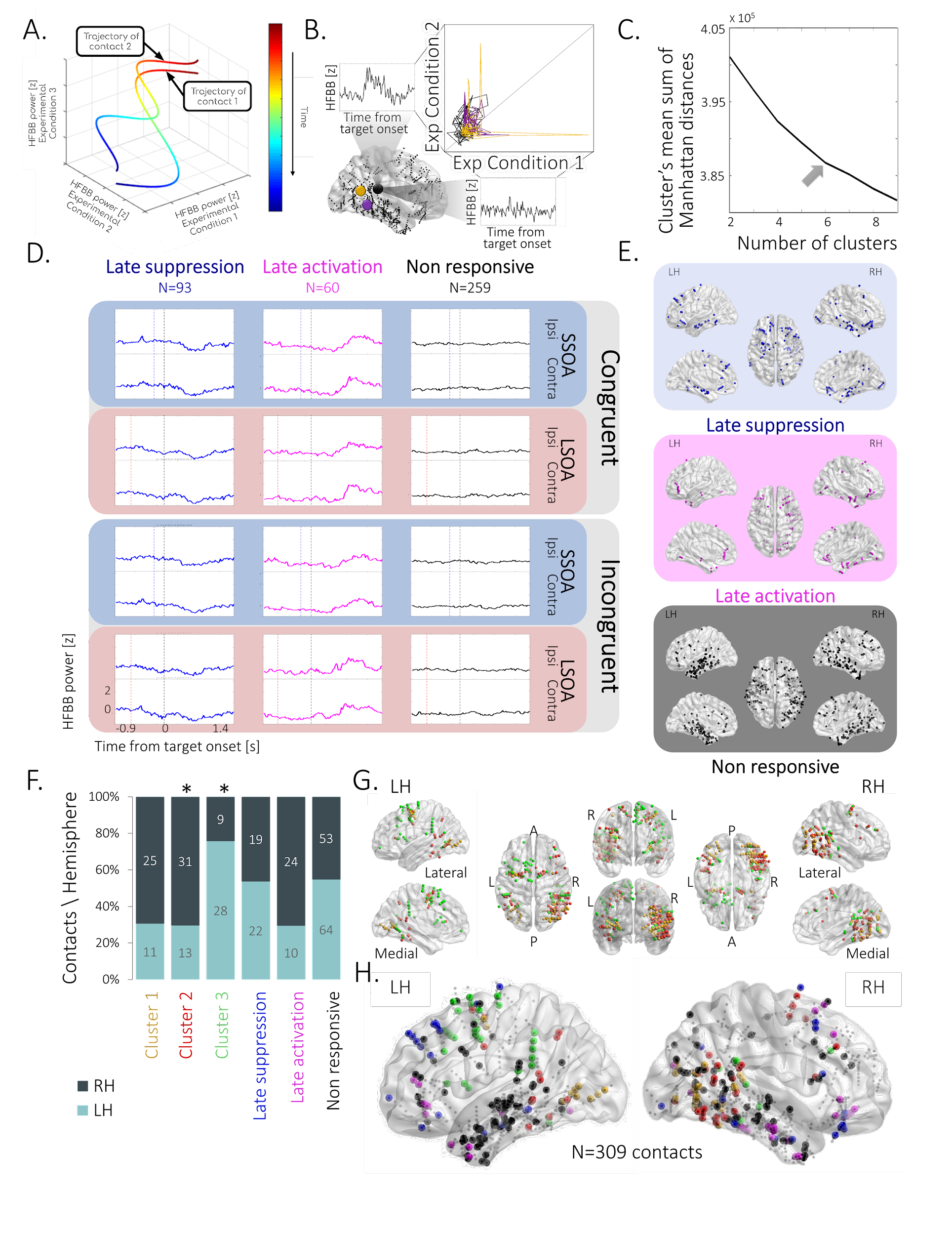

**Figure S2** – Clusters’ spatiotemporal profiles. (A) Illustration of neural activity trajectories of two contacts in a simplified 3-D condition space, color-coded by the temporal order of the sampled neural activity, progressing from red to blue. The trajectories represent the contacts’ neural activity measured as HFBB power in all three experimental conditions simultaneously. (B) A simplified example transformation of HFBB time series traces of a contact (black) in two experimental conditions into a neural activity trajectory in a 2-D condition space, represented along with the trajectories of two other contacts (yellow & purple). Contact locations in the brain are depicted in the lower left inset (black, yellow & purple circles). (C) Elbow method - mean sum of Manhattan distances between each contact trajectory and its assigned cluster trajectory for 2-9 clusters’ solutions; maximal elbow (grey arrow) at k=6. (D) Prototypical target-locked activity profiles (Trimmed-mean) of Late suppression (blue), Late activation (magenta), and Non-responsive (black) clusters, across the 8 conditions (Congruent / Incongruent X short-SOA / long-SOA X Ipsilateral target (Ipsi) / contralateral target (Contra)), not included in the main analysis. Dashed vertical lines represent target (black), and short-SOA (blue) and long-SOA (red) cues onsets. (E) Localization of contacts of Late suppression (blue), Late activation (magenta) and Non-responsive (black) clusters, not included in the main analysis. Note that the Non-responsive cluster contained contacts with potentially idiosyncratic or induced (as opposed to evoked) responses, averaged out. Dots represent contacts’ localizations (mean coordinates of the contacts composing each bipolar montage) depicted in normalized space (MNI152) in dorsal (middle), lateral (top) and medial (bottom) views in the right (RH; right) and left hemispheres (LH; left). (F) Hemispheric asymmetry of cluster distribution in regions with similar coverage (Contingency tables analysis, *χ²*_(5)_=29.09, *p*<0.001, *n*=309 independent contacts). For each cluster, the bar’s color proportion represents % contacts per hemisphere with raw contact numbers per hemisphere. Cluster 2 (red) is right-lateralized and Cluster 3 (green) is left-lateralized (post hoc binomial tests, *p*=0.01 and *p*=0.003). (G) Localization of contacts of clusters 1, 2 & 3 (yellow, red & green, correspondingly) from different views. (H) Localization of clusters’ contacts in similarly covered regions (large dots color-coded according to (F); small dots denote recorded contacts not included in this analysis).

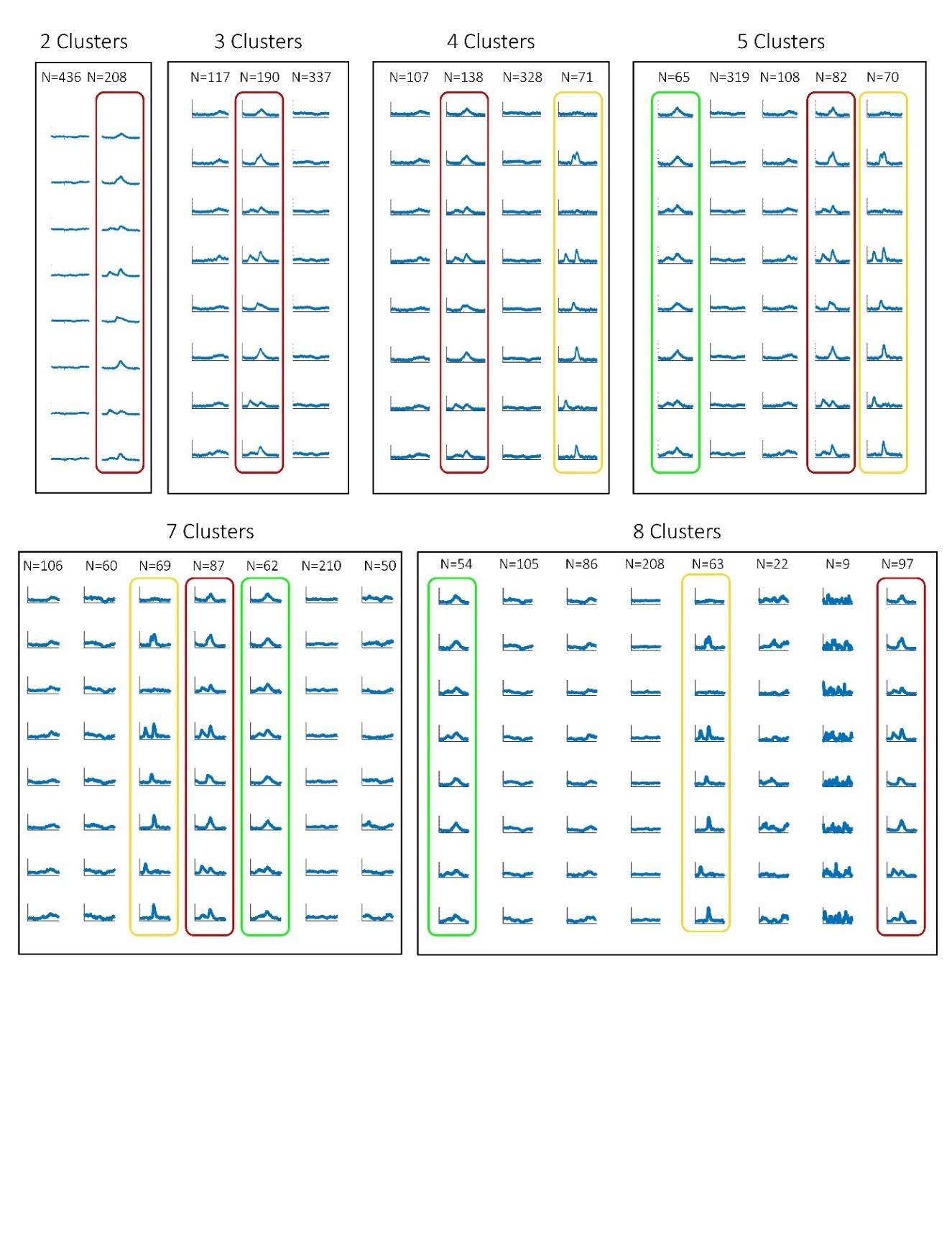

**Figure S3** – Trajectory k-means solution for different number of clusters. The three target-locked clusters analyzed in this study: Cluster 1 (yellow), Cluster 2 (red) and Cluster 3 (green) are present from 5-cluster solution onward, based on a contingency tables analysis showing a significant strong correspondence between each of the k-solutions and the 6-cluster solution (see Table S1 for details).

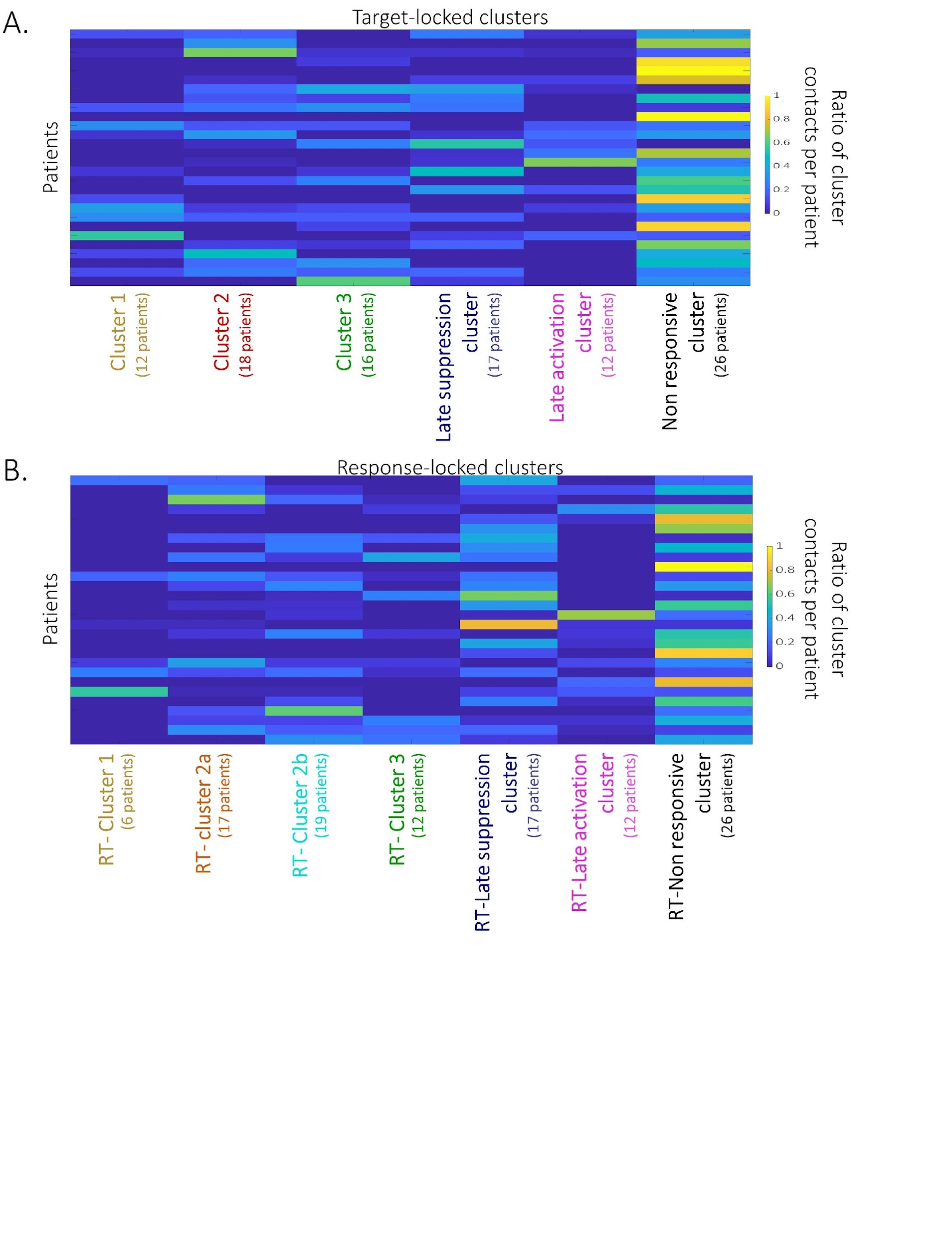

**Figure S4** – Distribution of the cluster contacts within participants. (A) The distribution of participants’ contributions to target-locked clusters. (B) The distribution of participants’ contributions to response-locked clusters. Each row represents one participant. Color code denotes the ratio of contacts in each cluster per participant.

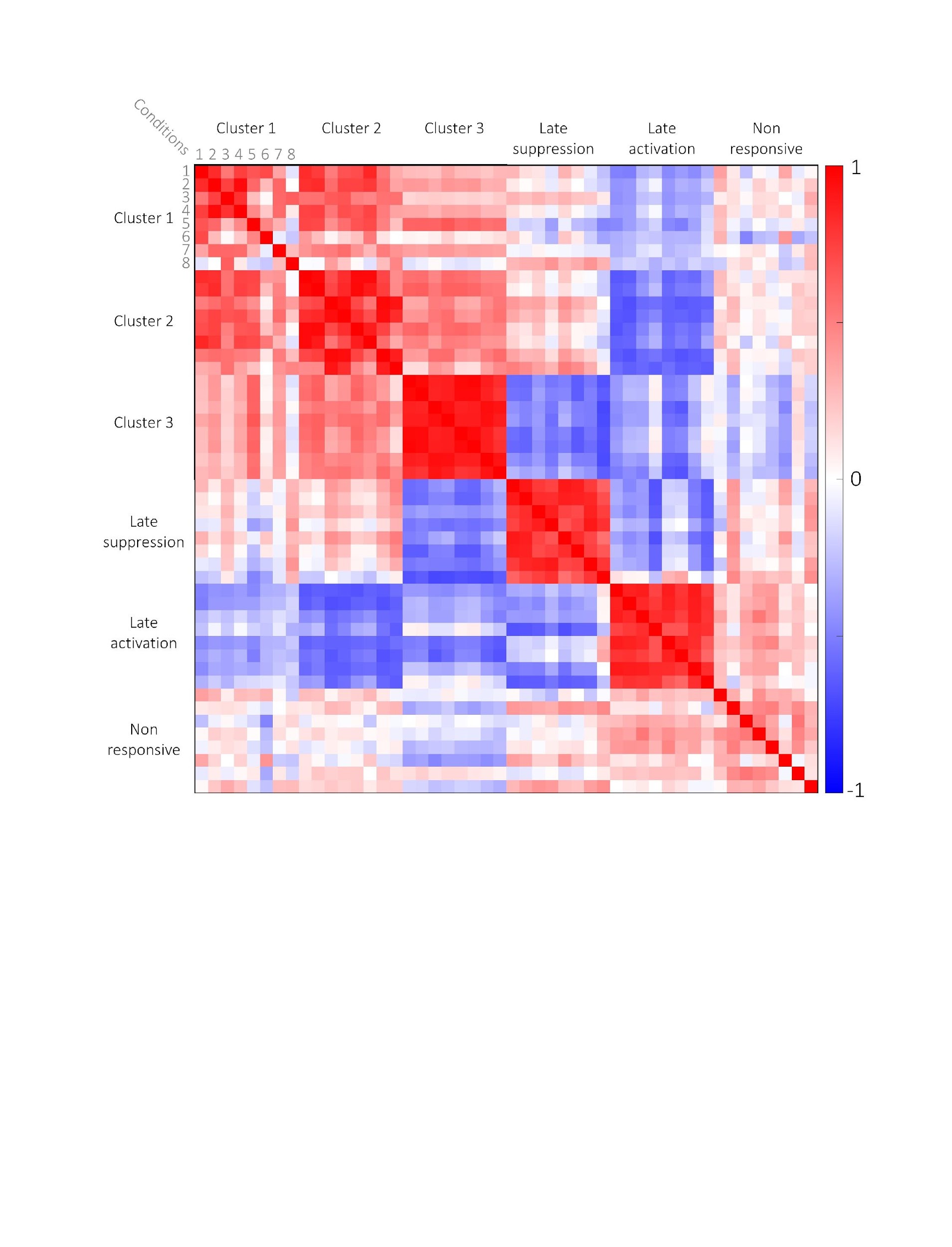

**Figure S5** – Clusters 1, 2 & 3 form a distinct group among all clusters. Pearson correlations between conditions’ centroid time-series across target-locked clusters reveal that the correlations of Clusters 1, 2 & 3 vary across experimental conditions within each cluster and positively correlate between these clusters. The correlation pattern within the three other clusters is more uniform and is negatively correlated across clusters. Color bar represents the r coefficient (negative correlation – blue; positive correlation-red); Numbers correspond to experimental conditions (1- Contralateral target short-SOA Congruent; 2- Contralateral target short-SOA Incongruent; 3-Contralateral target long-SOA Congruent; 4-Contralateral long-SOA Incongruent; 5-Ipsilateral target short-SOA Congruent; 6-Ipsilateral target short-SOA Incongruent; 7-Ipsilateral target long-SOA Congruent; 8-Ipsilateral target long-SOA Incongruent).

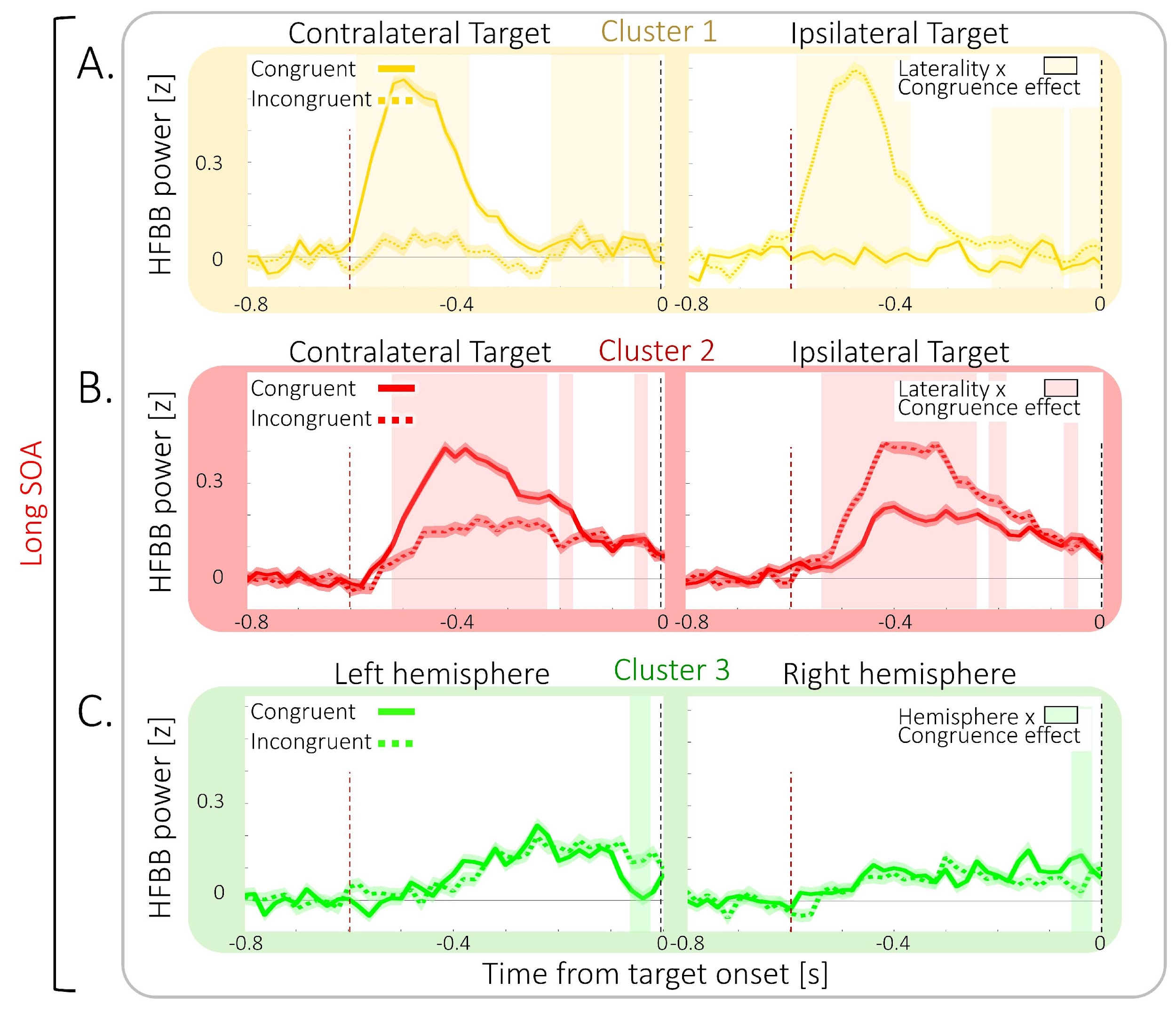

**Figure S6** – Congruence-related neural activity in the Cue time-window. Mean target-locked long-SOA activity pooled across all cluster contacts is depicted for Cluster 1 (yellow), Cluster 2 (red), and Cluster 3 (green). Full lines indicate Congruent trials and dashed lines represent Incongruent trials in the long-SOA condition. (A) In Cluster 1, a significant Laterality x Target-congruence effect was observed (yellow shaded area; time-resolved 3-way ANOVA; largest p=0.018), indicating it only responds to contralateral cues. (B) Cluster 2 exhibited stronger responses to contralateral cues than to ipsilateral ones, as evidenced by a significant Laterality x Target-congruence effect (shaded red areas; largest p=0.038). (C) Cluster 3 demonstrated a significant Hemisphere x Target-congruence effect (green shaded area; 3-way ANOVA, largest p=0.045). Shaded areas around traces represent standard error of the mean (SEM), dashed vertical lines indicate target onset (black) and Cue onset (red).

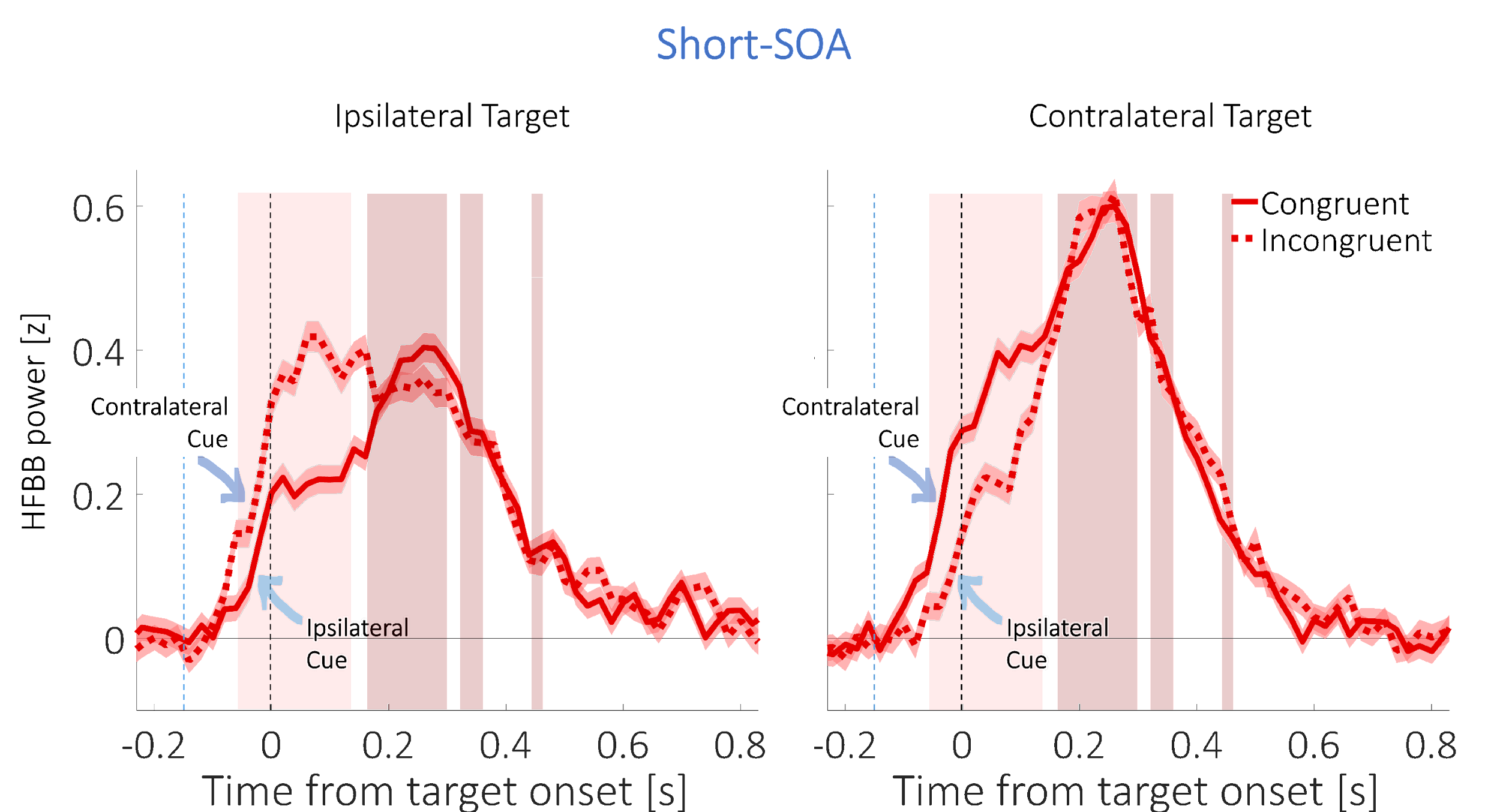

**Figure S7** – Exploratory analysis of short-SOA congruence-related neural activity in the target time-window in Cluster 2. Mean target-locked short-SOA activity in Cluster 2 (red) computed over trials pooled across all cluster contacts, for Congruent trials (full lines) and Incongruent trials (dashed lines), when targets were ipsilateral (left) or contralateral to the recording contact (right). Note, that when targets were ipsilateral, Incongruent cues were contralateral (dark blue arrow), and Congruent cues were ipsilateral (light blue arrow), and conversely for contralateral targets. Responses were stronger to contralateral cues and targets than to ipsilateral ones, as shown by a 3-way ANOVA significant Target-side x Congruence effect (shaded light red areas; -60-140ms post target onset; largest *p*=0.022) and a main Target-side effect (shaded dark red areas; 160-300ms; 320-360ms; 440-460ms post Target onset; largest *p*=0.012). Shaded areas around traces depict SEM; Dashed vertical lines represent target onset (black) and Cue onset (blue).

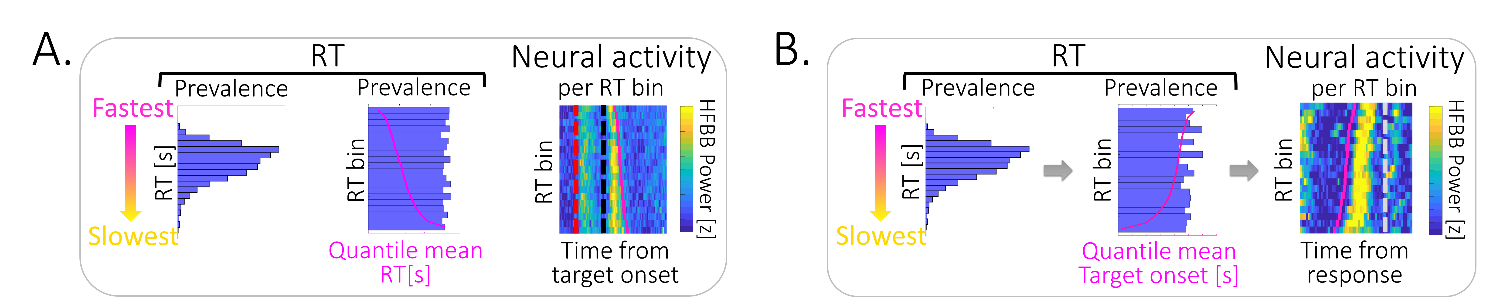

**Figure S8** - (A) Computation of neural activity across RT bins: Within each cluster, the trial distribution of RTs across all conditions (left) was divided into 20 quantiles (RT bins; middle), ordered by mean RT (magenta line). The quantile’s mean target-locked neural activity pooled across contacts of each cluster was computed (right; Vertical dashed lines denote cue (red) & target (black) onset; magenta line represent mean RT). (B) Computation of neural response-locked activity across RT bins: Within each cluster, the trial distribution of RTs in each condition (left) was divided into 20 quantiles (RT bins; middle), ordered by mean RT, here corresponding to target onset time (magenta line). The mean Response-locked neural activity across all cluster contacts for each quantile was computed (right; Vertical grey dashed line denote RT (black) onset; magenta line represent mean target onset time).

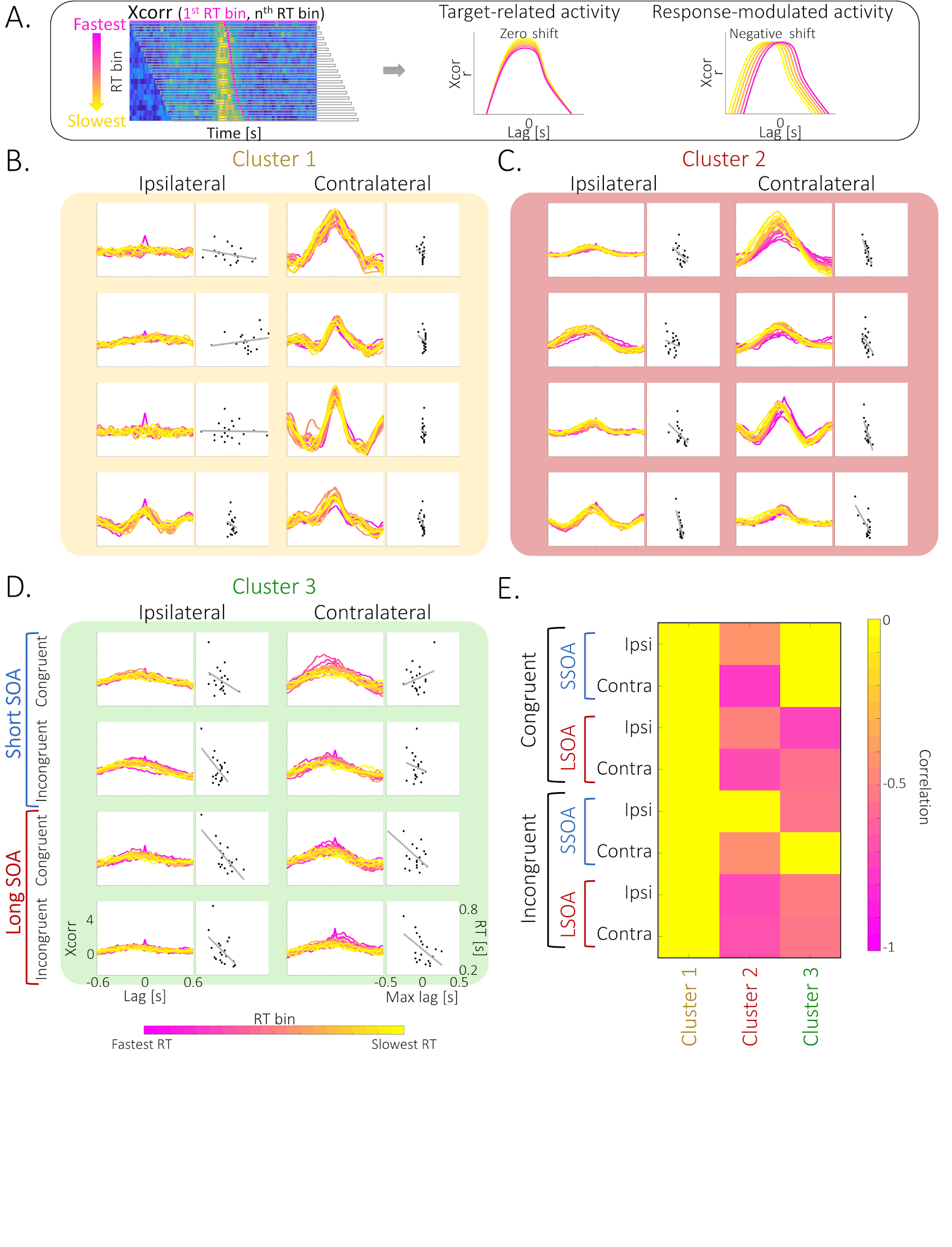

**Figure S9** – Clusters’ neural target-locked activity timing correlates with behavior. (A) Schematic illustration of the procedure for computing the cross-correlation (Xcorr) of neural activity across RT bins: Cross-correlation between target-locked activity at the fastest RT bin and all subsequent bins was computed (left). If cluster activity is target-associated, maximal cross-correlation will be centered on target onset, resulting in a zero lag shift across all RT bins (middle). If cluster activity is response-associated, maximal cross-correlation will follow the RT, resulting in a negative shift of cross-correlation lag (right). (B)-(D). Cross-correlogram of neural activity at different RT bins (pink- fastest RT; yellow - slowest RT) as a function of cross-correlation lag (left columns) and Pearson correlation (grey line) between maximal cross-correlation lags (Max lag) and bin’s mean RTs (right columns), across the 8 conditions (Congruent / Incongruent X short-SOA / long-SOA X Ipsilateral target / contralateral target) in Cluster 1 (yellow), Cluster 2 (red) and Cluster 3 (green). (B) Cluster 1 activity is target-associated: Cross-correlation plots are centered on zero, especially for contralateral targets. (C) Activity in Cluster 2 is response-associated: Cross-correlation plots show a negative shifted lag that is generally correlated with RT. (D) Cluster 3 activity is response-associated: Cross-correlation plots show a negative shifted lag, correlated with RT under certain conditions. (E) Significant negative correlation between cross-correlation maximal lag and bin mean RT in Clusters 2 & 3: significant (p<0.05) negative correlations were found only in these two clusters.

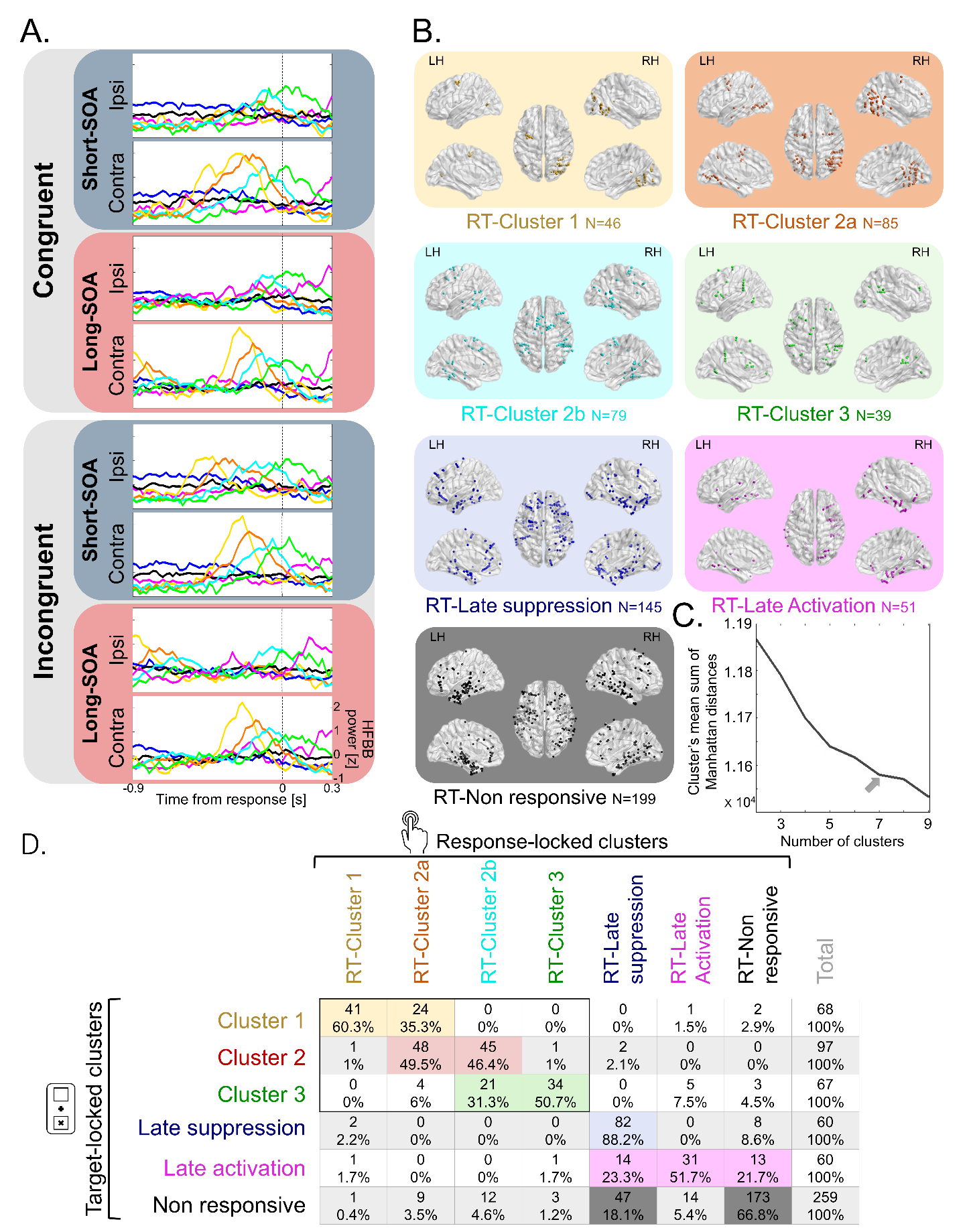

**Figure S10** – Spatiotemporal profiles of Response-locked clusters. (A) Trimmed-mean Response-locked activity profiles of the seven contact clusters across the 8 conditions (Congruent / Incongruent X short-SOA / long-SOA X Ipsilateral target / contralateral target): RT-Cluster 1 (yellow); RT-Cluster 2a (orange); RT- Cluster 2b (turquoise); RT-Cluster 3 (green); RT-Late suppression cluster (blue); RT-Late activation cluster (magenta); RT-Non responsive cluster (black). Dashed vertical line represents motor response time. (B) Response-locked clusters’ spatial location. Illustration of the localization of the contacts composing each cluster (colors as in A). For each cluster, dots represent contacts’ localization, computed as the mean coordinates of the two contacts composing each contact’s bipolar montage, depicted in normalized space (MNI152) in dorsal (middle), lateral (top) and medial (bottom) views in the right hemisphere (RH) and the left hemisphere (LH). (C) Elbow method. Mean sum of Manhattan distances between each contact trajectory and its assigned cluster trajectory for 2-9 clusters’ solution. Maximal elbow (grey arrow) is observed at 7-cluster solution. (D) Mapping between target-locked and response-locked clusters. The distribution of target-locked clusters’ contacts (rows; number of contacts & % within row) across the different response-locked clusters (columns) was significantly different than chance (Contingency tables analysis, *p*<0.001; Contingency coefficient =0.83, N=259 contacts).

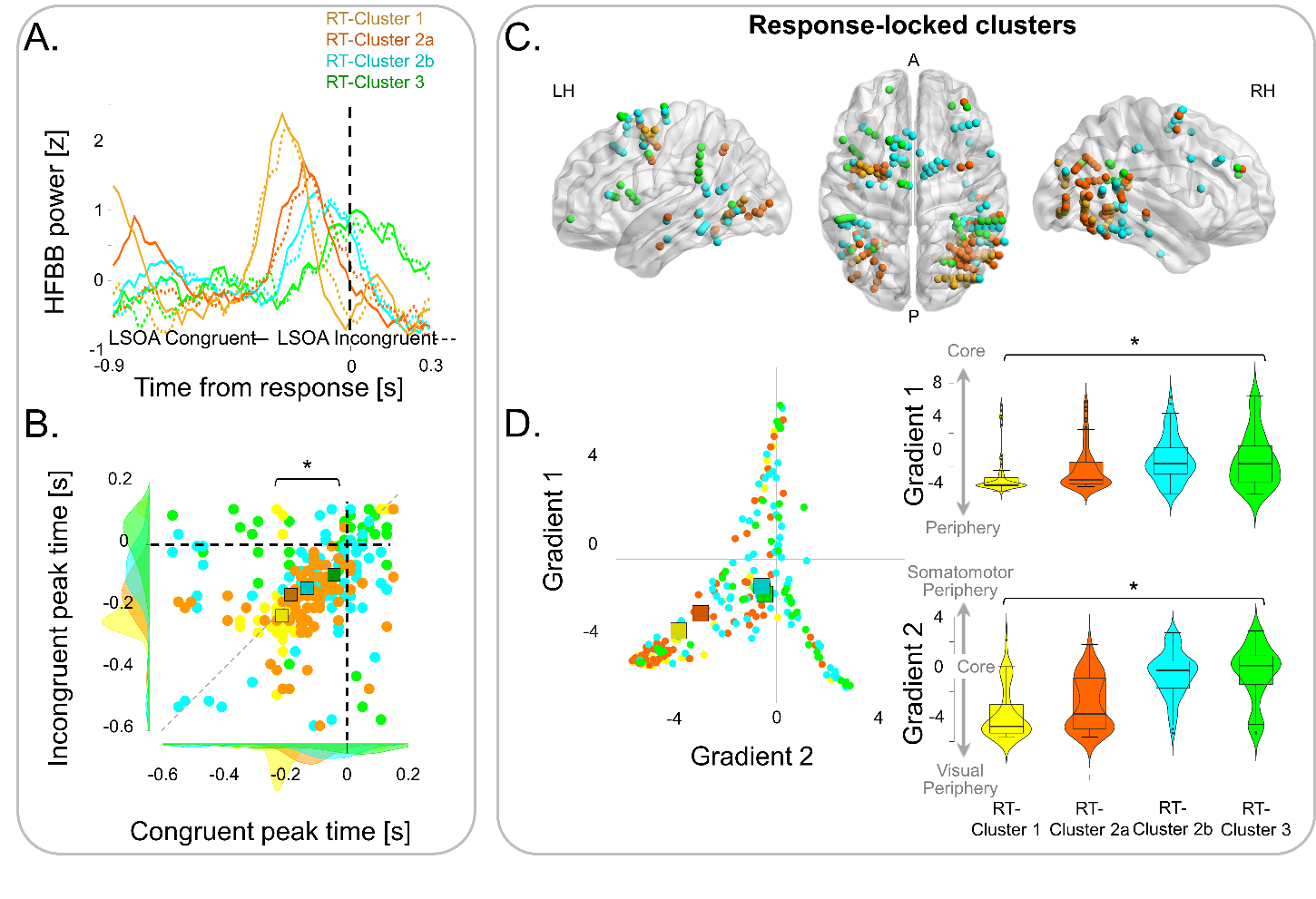

**Figure S11** – Response-locked clusters exhibit a spatiotemporal gradient. (A) Temporal gradient of activity in Response-locked clusters: Trimmed-mean Response-locked activity of RT-Cluster 1, RT-Cluster 2a, RT-Cluster 2b and Cluster 3. Black dashed line depicts RT. (B) Scatter plot of peak times of mean Response-locked activity of the contacts of RT-Cluster 1 (yellow circles), RT-Cluster 2a (orange circles), RT-Cluster 2b (turquoise circles) and RT-Cluster 3 (green circles) in the Congruent (x axis) and Incongruent (y-axis) long-SOA conditions, showing a significant temporal gradient (Mixed Anova: Cluster main effect *F*(3,245)=12.57, *p*<0.001, *η^2^*=0.086; linear polynomial contrast: p≤0.001). Squares represent mean peak time; Dotted grey line denotes the equity line; Shaded areas represent peak time distributions. (C) Core-Periphery gradient: Clusters’ anatomical localization follows Core-Periphery gradients (Margulies et al., 2016), where RT-Cluster 1’s contacts are the most peripheral and RT-Cluster 3’s contacts are closest to core regions. (D) Left: Scatter plot of contacts localization along core-periphery gradients (RT-Cluster 1 - yellow circles, n=46 independent contacts; RT-Cluster 2a - orange circles, n=85 independent contacts; RT-Cluster 2b – turquoise circles, n=79 independent contacts; RT-Cluster 3 - green circles, n=39 independent contacts). Top & bottom right: Violin plots of contacts localization along Core-Periphery gradients for RT-Cluster 1 (yellow), RT-Cluster 2a (orange), RT-Cluster 2b (turquoise) and RT-Cluster 3 (green) clusters, showing a significant core-periphery gradient (Gradient 1: 1-way ANOVA, *p*=0.001, *η^2^*=0.06; linear polynomial contrast: p≤0.001; Gradient 2: 1-way ANOVA, p<0.001, *η^2^*=0.32; linear polynomial contrast: p≤0.001). The box centerlines depict the medians, the bounds of the box depict the 75%/25% quartiles and the whiskers depict the top & bottom 25% percentiles.

### Supplementary Tables

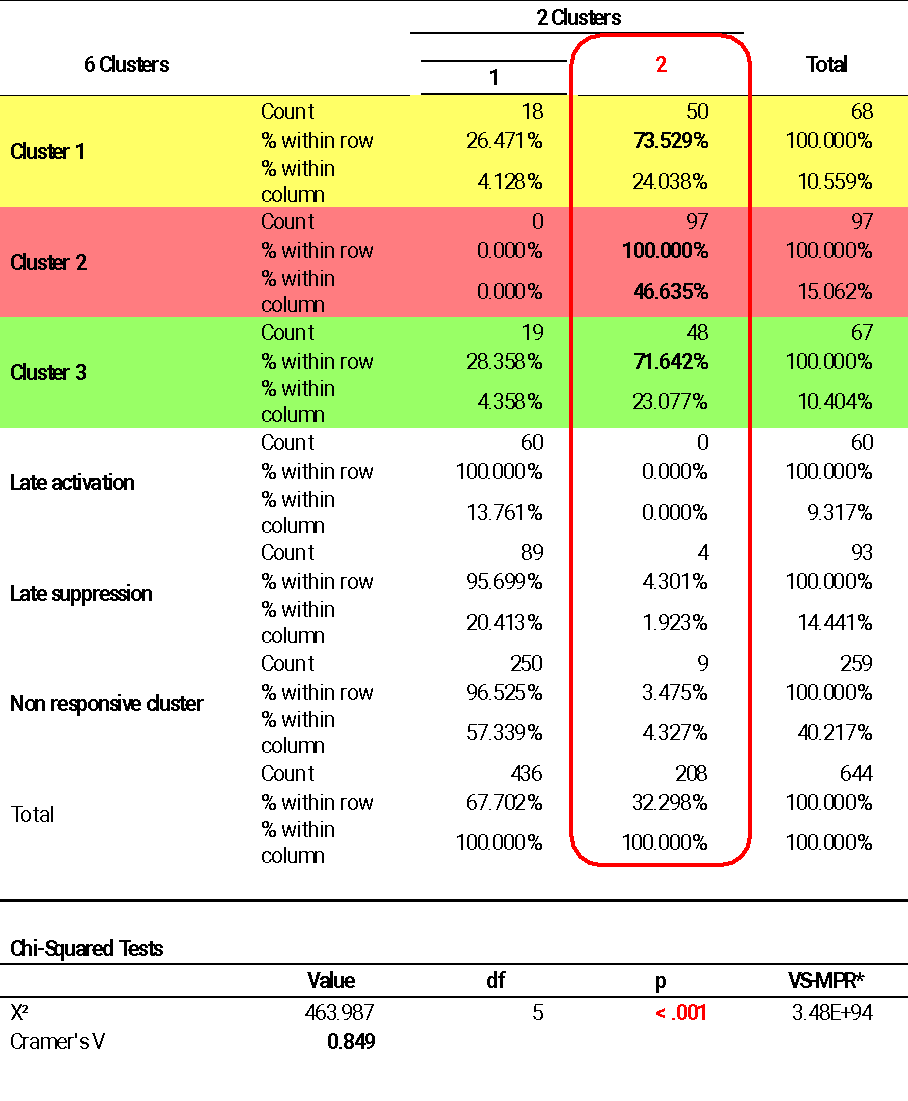

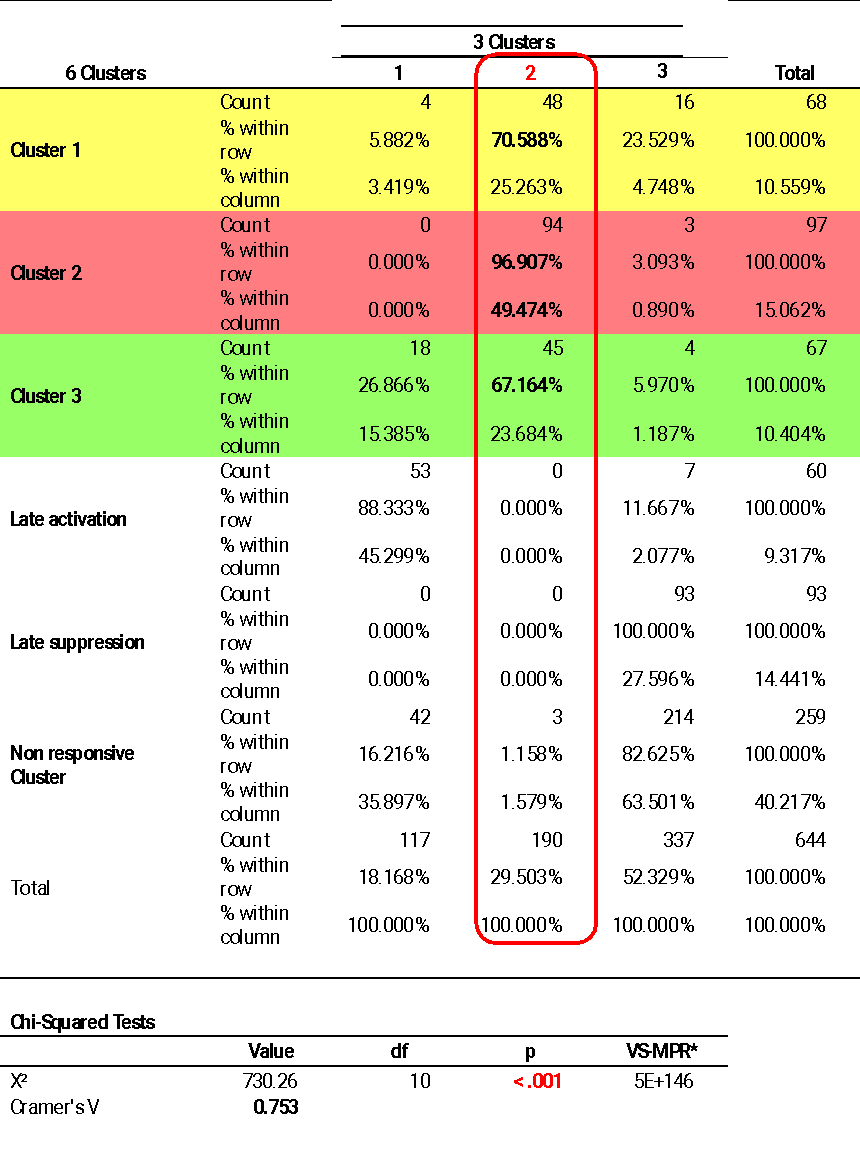

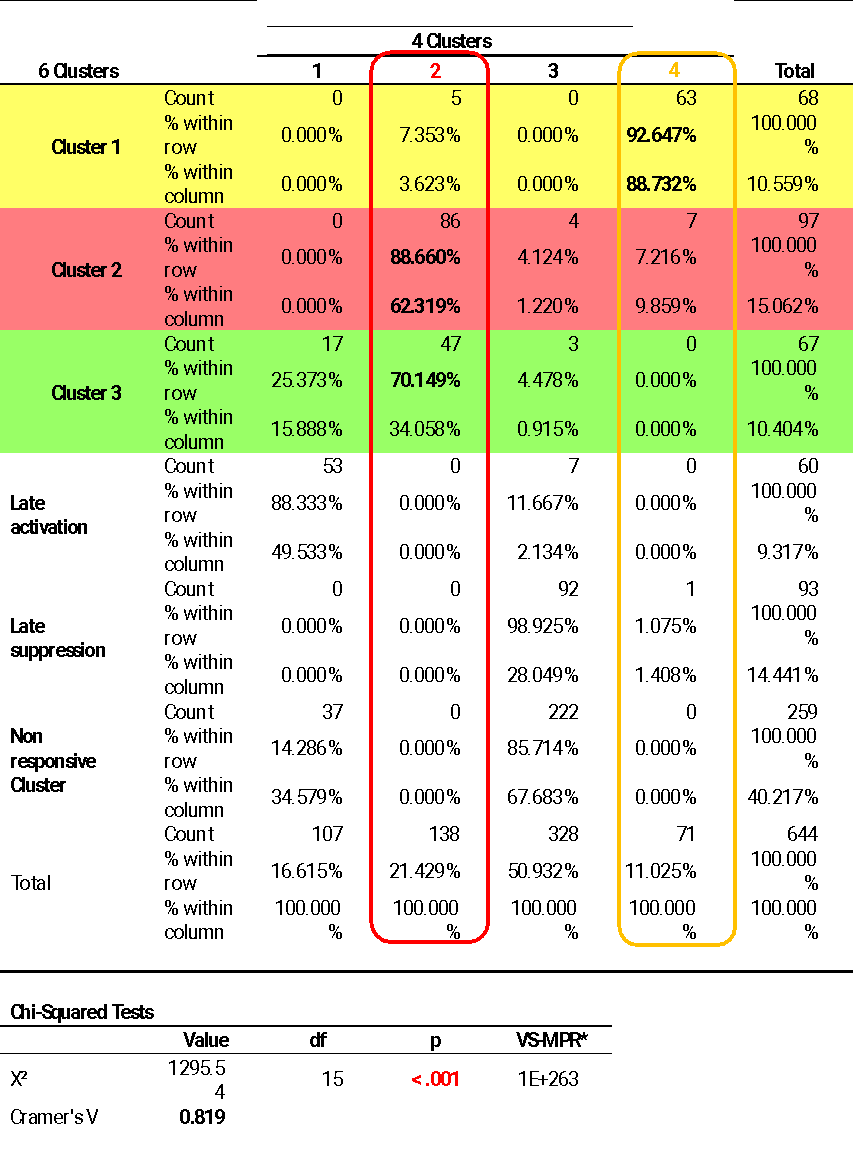

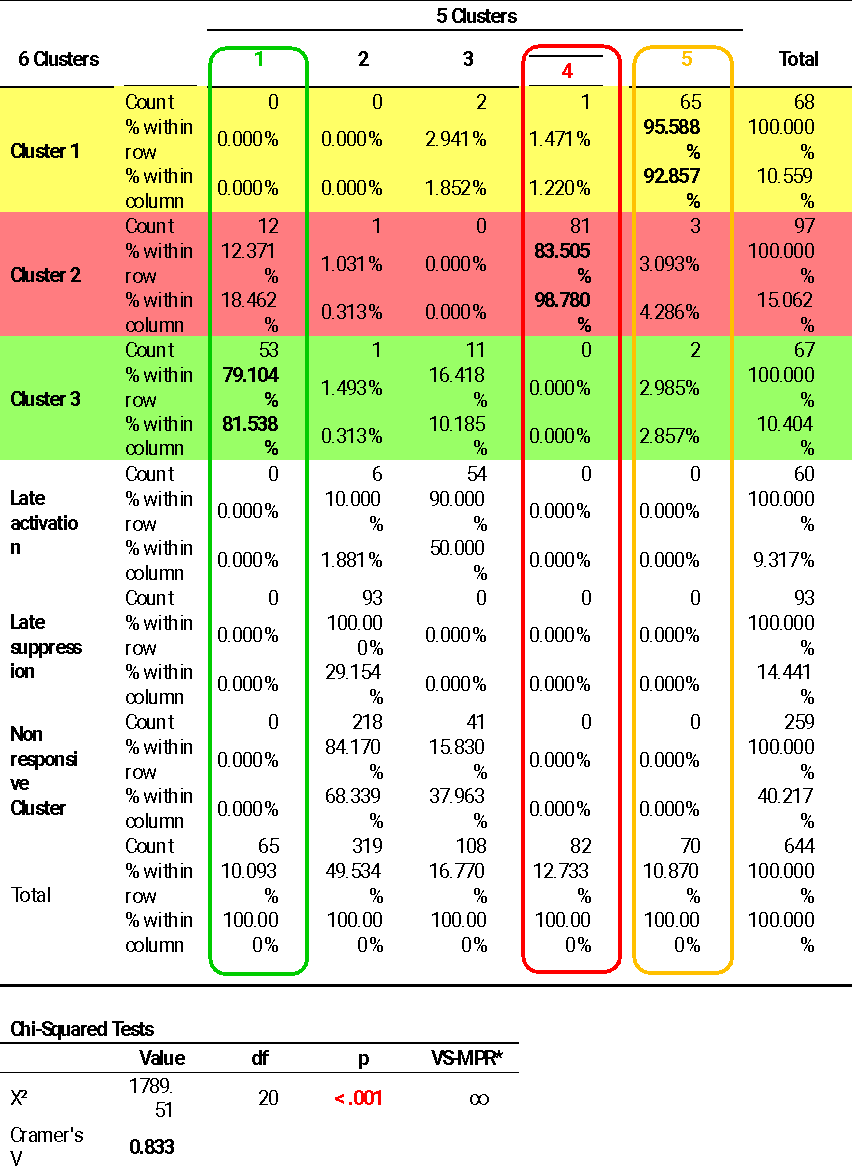

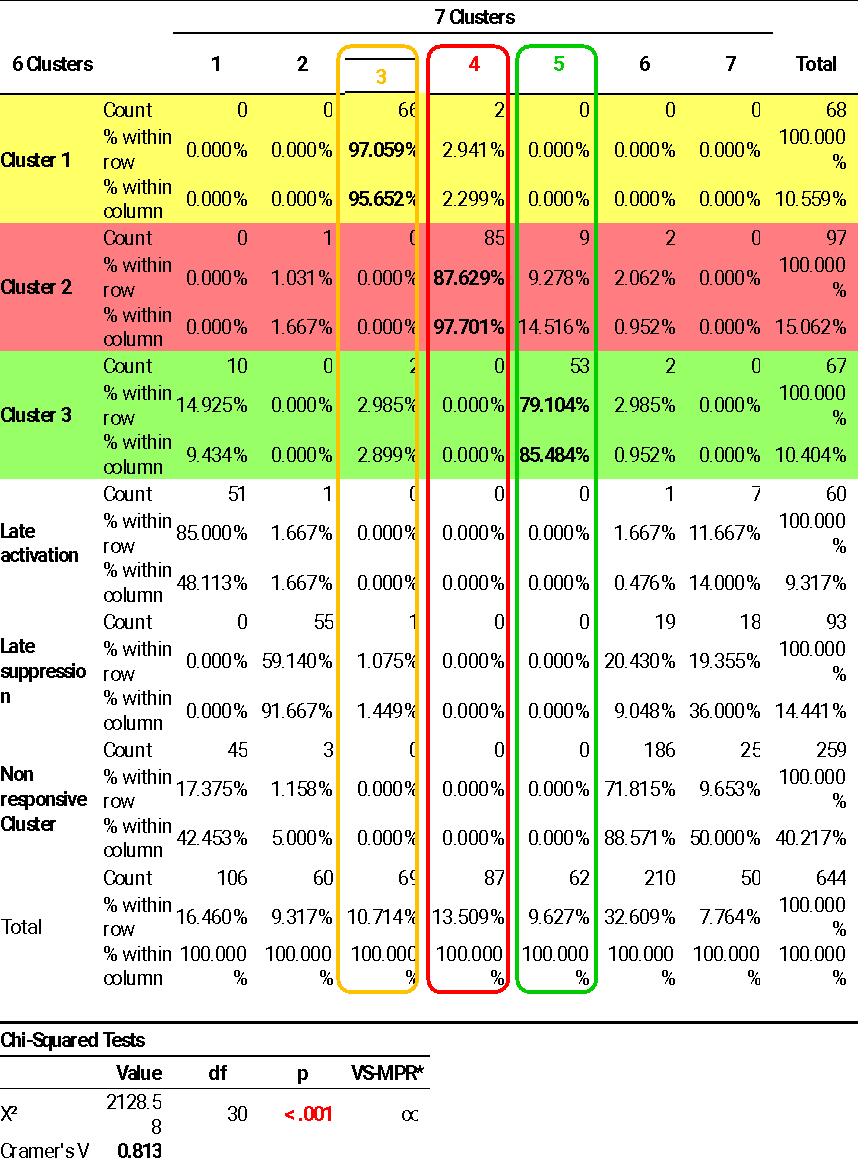

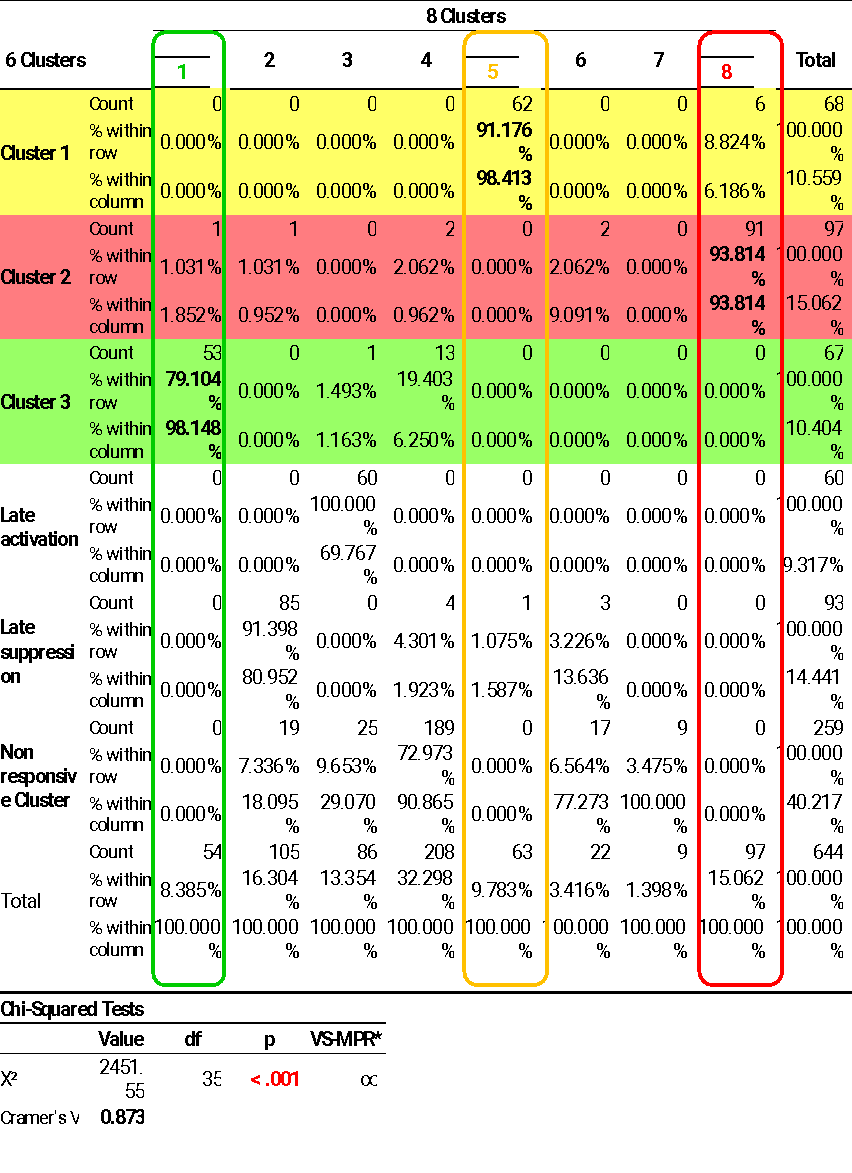
 **Table S1** – Cluster stability across 2-8 *k*-cluster solutions. Strong significant correspondence (Contingency tables analyses, all *p*<0.001, all Cramer’s *V*≥0.75) between the assignments of contacts to clusters in the 6-cluster solution and the other k-cluster solutions (from *k*=2 in the top table to *k*=8 in the bottom table). The Contingency tables show the distribution of contacts belonging to each of the three further analyzed clusters (Cluster 1- yellow, Cluster 2 – red, Cluster 3 – green) in each of the clusters of the other *k* solutions (% within row), and the composition of each of the other solutions’ clusters (% within column). A *k-*solution cluster was marked as stable (colored frame) if the main group of contacts composing it could be mapped to one of the three further analyzed clusters, which in turn shared most of its contacts with that cluster.

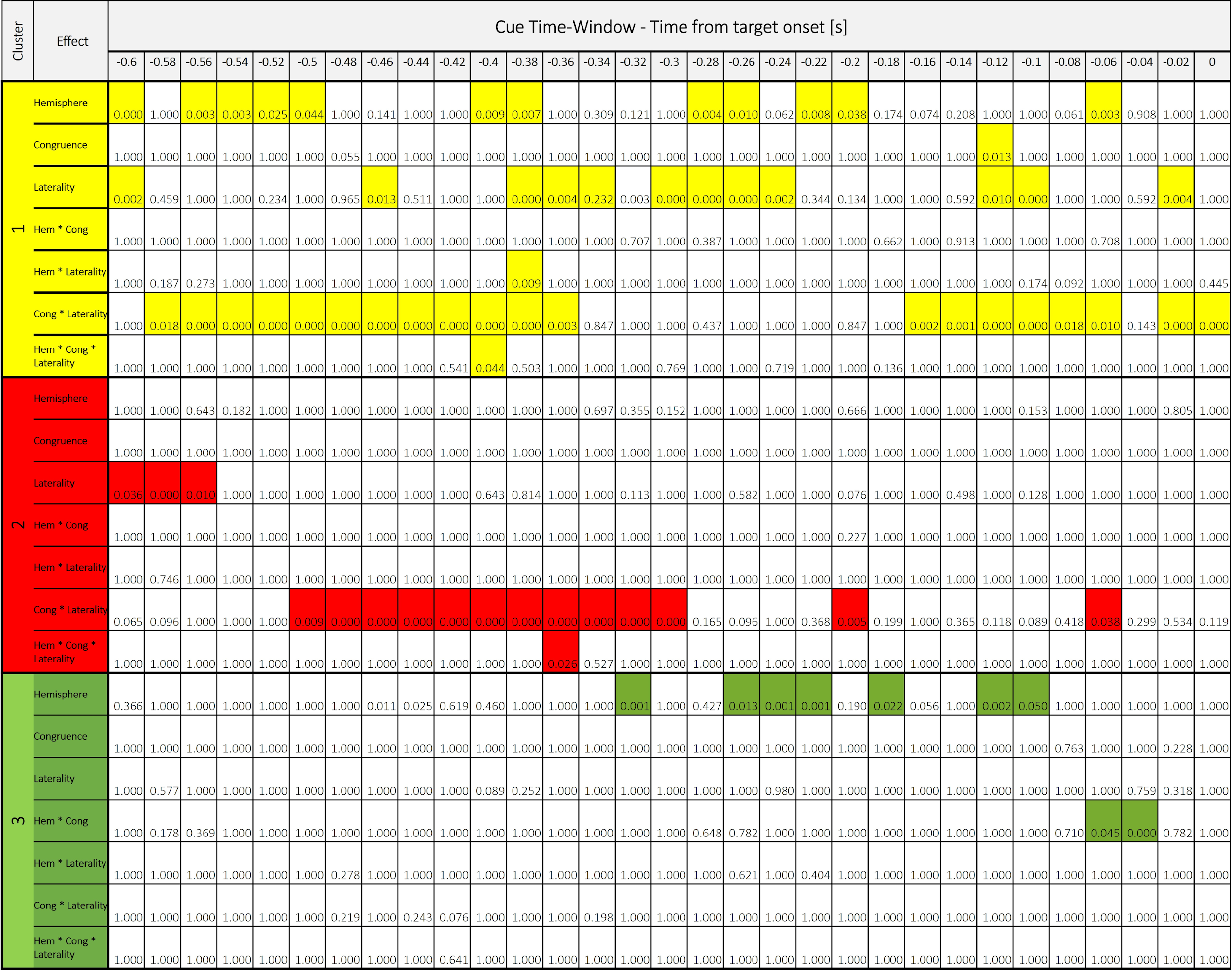

**Table S2** – IOR-related neural activity in the cue time-window. Holm corrected p-values for the 3-way ANOVA testing the effects of Congruence, Hemisphere and Target-side on the HFBB signal in the long-SOA condition in Cluster 1 (yellow), Cluster 2 (red) and Cluster 3 (green). Significant effects in shaded color.

**Table S3** – IOR-related neural activity in the Target time-window. Holm corrected p-values for the 3-way ANOVA testing the effects of Congruence, Hemisphere and Target-side on the HFBB signal in the long-SOA condition in Cluster 1 (yellow), Cluster 2 (red) and Cluster 3 (green). Significant effects in shaded color.
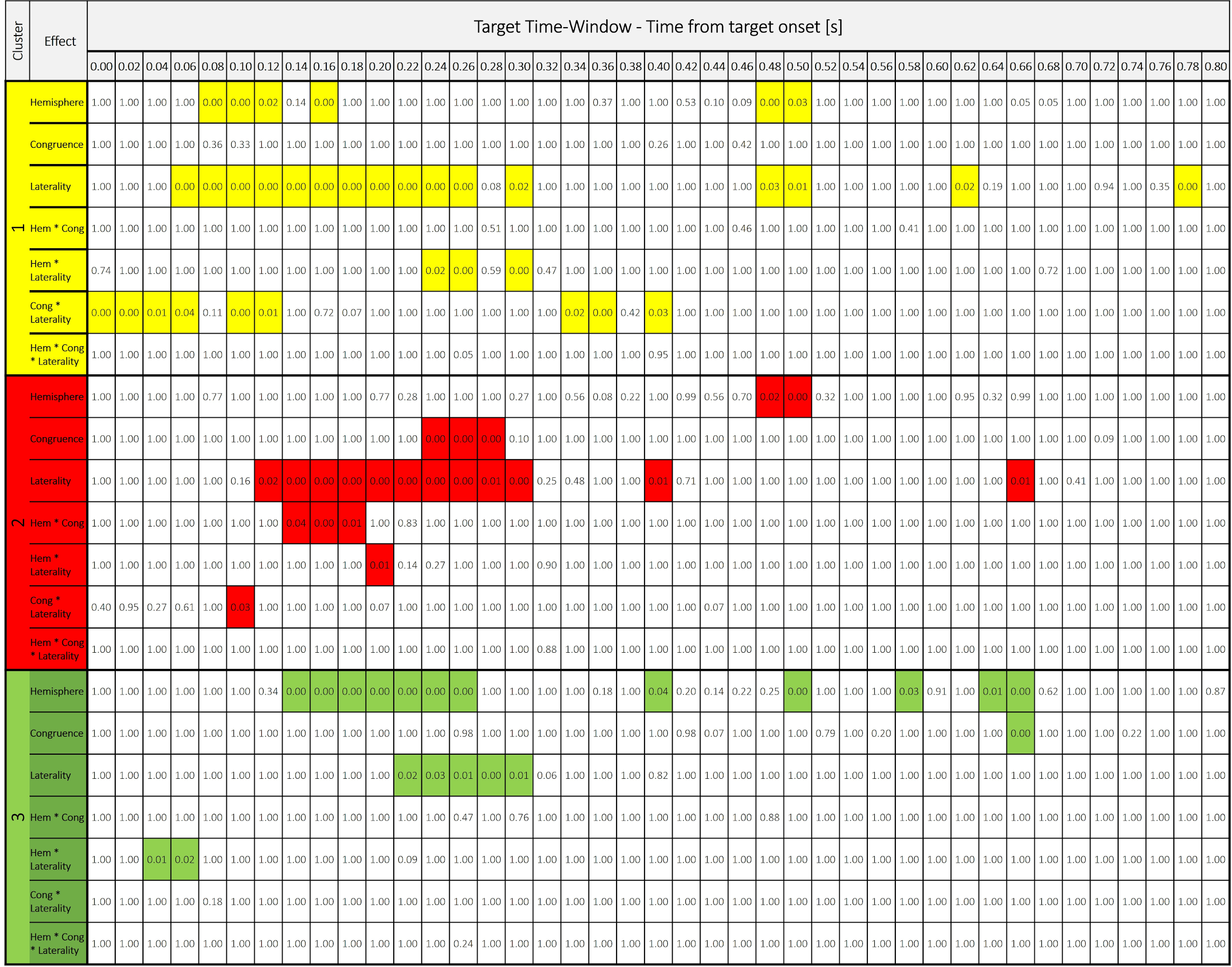

### Supplementary Methods

Cluster hemispheric lateralization

The hemispheric lateralization of the clusters was tested on a subgroup of contacts localized in cortical volumes that were sampled in both hemispheres. This was done to overcome the confound of unequal coverage within the hemispheres. To identify similarly-covered contacts, a 3mm radius sphere (corresponding to the assumed volume recorded by iEEG contacts ^2^) was fit around each contact using SPM12 ^3^, and the overlap between each of the spheres and the entire covered volume in the other hemisphere was calculated. The cluster-distribution of the 309 resulting contacts (148 in the left hemisphere and 161 in the right hemisphere) across the hemispheres was compared using a contingency table analysis in JASP ^4^, and post hoc binomial test with Holm correction were conducted to identify the clusters with significant hemispheric lateralization.

*Cross-correlation of target-locked and response-locked RT-bins*

This analysis intended to explore the link between target-locked neural activity and RT. We sorted in each cluster all the trials pooled over the conditions according to their RT. We then binned them into 20 quantiles (Fig. S3A). In each cluster, a cross-correlation between target-locked activity at the fastest RT bin and all subsequent bins was computed for each experimental condition within a maximal lag of ±600 ms. If cluster activity is target-related, maximal cross-correlation will be centered on target onset, resulting in a zero shift across all RT bins. If cluster activity is response-modulated, maximal cross-correlation will follow the RT, resulting in a negative shift of cross-correlation lags. Next, in order to test whether the maximal lag corresponded to the actual RT (expecting significant negative correlation coefficients for RT-modulated activity), a Pearson correlation between the maximal cross-correlation lag and each bin’s mean RT was computed for each condition. Cross-correlation between response-locked activity at the fastest RT bin and all subsequent bins was similarly computed. Notably, if cluster response-locked activity is visually modulated, maximal cross-correlation will follow the RT (here marking quantile’s mean target-onset time), resulting in a positive shift of the maximal cross-correlation lag. If cluster activity is only response-associated, maximal cross-correlation will be centered on target onset, resulting in a zero shift across all RT bins.

*Theta-phase dependence of neural activity*

The instantaneous theta (4-8Hz) phase was extracted from the raw unfiltered data using a hilbert transform. The phase angle at the onset of the target stimulus was compared between conditions with different SOAs and congruence level using a mixed ANOVA with repeated-measures factors of SOA and Congruence, supplemented by a between-subjects factor of Cluster to test if the theta phase effect could arise differentially across different contact clusters.
